# Probabilistic variable-length segmentation of protein sequences for discriminative motif discovery (DiMotif) and sequence embedding (ProtVecX)

**DOI:** 10.1101/345843

**Authors:** Ehsaneddin Asgari, Alice McHardy, Mohammad R.K. Mofrad

## Abstract

In this paper, we present peptide-pair encoding (PPE), a general-purpose probabilistic segmentation of protein sequences into commonly occurring variable-length sub-sequences. The idea of PPE segmentation is inspired by the byte-pair encoding (BPE) text compression algorithm, which has recently gained popularity in subword neural machine translation. We modify this algorithm by adding a sampling framework allowing for multiple ways of segmenting a sequence. PPE segmentation steps can be learned over a large set of protein sequences (Swiss-Prot) or even a domain-specific dataset and then applied to a set of unseen sequences. This representation can be widely used as the input to any downstream machine learning tasks in protein bioinformatics. In particular, here, we introduce this representation through protein motif discovery and protein sequence embedding. (i) DiMotif: we present DiMotif as an alignment-free discriminative motif discovery method and evaluate the method for finding protein motifs in three different settings: (1) comparison of DiMotif with two existing approaches on 20 distinct motif discovery problems which are experimentally verified, (2) classification-based approach for the motifs extracted for integrins, integrin-binding proteins, and biofilm formation, and (3) in sequence pattern searching for nuclear localization signal. The DiMotif, in general, obtained high recall scores, while having a comparable F1 score with other methods in the discovery of experimentally verified motifs. Having high recall suggests that the DiMotif can be used for short-list creation for further experimental investigations on motifs. In the classification-based evaluation, the extracted motifs could reliably detect the integrins, integrin-binding, and biofilm formation-related proteins on a reserved set of sequences with high F1 scores. (ii) ProtVecX: we extend k-mer based protein vector (ProtVec) embedding to variable-length protein embedding using PPE sub-sequences. We show that the new method of embedding can marginally outperform ProtVec in enzyme prediction as well as toxin prediction tasks. In addition, we conclude that the embeddings are beneficial in protein classification tasks when they are combined with raw k-mer features.

**Availability:** Implementations of our method will be available under the Apache 2 licence at http://llp.berkeley.edu/dimotif and http://llp.berkeley.edu/protvecx.

## 1 Introduction

Bioinformatics and natural language processing (NLP) are research areas that have greatly benefited from each other since their beginnings and there have been always methodological exchanges between them. Levenshtein distance [1] and Smith–Waterman [2] algorithms for calculating string or sequence distances, the use of formal languages for expressing biological sequences [3, 4], training language model-based embeddings for biological sequences [5], and using state-of-the-art neural named entity recognition architecture [6] for secondary structure prediction [7] are some instances of such influences. Similar to the complex syntax and semantic structures of natural languages, certain biophysical and biochemical grammars dictate the formation of biological sequences. This assumption has motivated a line of research in bioinformatics to develop and adopt language processing methods to gain a deeper understanding of how functions and information are encoded within biological sequences [4, 5, 8]. However, one of the apparent differences between biological sequences and many natural languages is that biological sequences (DNA, RNA, and proteins) often do not contain clear segmentation boundaries, unlike the existence of tokenizable words in many natural languages. This uncertainty in the segmentation of sequences has made overlapping k-mers one of the most popular representations in machine learning for all areas of bioinformatics research, including proteomics [5, 9], genomics [10, 11], epigenomics [12, 13], and metagenomics [14, 15]. However, it is unrealistic to assume that fixed-length k-mers are units of biological sequences and that more meaningful units need to be introduced. This means that although choosing a fixed k value for sequence k-mers simplifies the problem of segmentation, it is an unrealistic assumption to assume that all important part of the sequences have the same length and we need to relax this assumption.

Although in some sequence-labeling tasks (e.g. secondary structure prediction or binding site prediction) sequences are implicitly divided into variable-length segments as the final output, methods to segment sequences into variable-length meaningful units as inputs of downstream machine learning tasks are needed. We recently proposed nucleotide pair encoding for phenotype and biomarker detection in 16S rRNA data [16], which is extended in this work for protein informatics.

Here, we propose a segmentation approach for dividing protein sequences into frequent variable-length sub-sequences, called peptide-pair encoding (PPE). We took the idea of PPE from byte pair encoding (BPE) algorithm, which is a text compression algorithm introduced in 1994 [17] that has been also used for compressed pattern matching in genomics [18]. Recently, BPE became a popular word segmentation method in machine translation in NLP for vocabulary size reduction, which also allows for open-vocabulary neural machine translation [19]. In contrast to the use of BPE in NLP for vocabulary size reduction, we used this idea to increase the size of symbols from 20 amino acids to a large set of variable-length frequent sub-sequences, which are potentially meaningful in bioinformatics tasks. In addition, as a modification to the original algorithm, we propose a probabilistic segmentation in a sampling framework allowing for multiple ways of segmenting a sequence into sub-sequences. In particular, we explore the use of PPE for protein sequence motif discovery as well as training embeddings for protein sequences.

### *De novo* motif discovery

Protein short linear motif (SLiM) sequences are short sub-sequences of usually 3 to 20 amino acids that are presumed to have important biological functions; examples of such patterns are cleavage sites, degradation sites, docking sites, ligand binding sites, etc [20, 21]. Various computational methods have been proposed for the discovery of protein motifs using protein sequence information. Motif discovery is distinguished from searching for already known motifs in a set of unseen sequences (e.g., SLiMSearch [22]). Motif discovery can be done either in a discriminative or a non-discriminative manner. Most of the existing methods are framed as non-discriminative methods, i.e. finding overrepresented protein sub-sequences in a set of sequences of similar phenotype (positive sequences). Examples of non-discriminative methods are SLiMFinder [23] (regular expression based approach), GLAM2 [24] (simulated annealing algorithm for alignments of SLiMs), MEME [25] (mixture model fitting by expectation-maximization), HH-MOTiF [26] (Hidden Markov Model (HMM) model based approach on multiple sequence alignment). Since other randomly conserved patterns may also exist in the positive sequences, reducing the false positive rate is a challenge for motif discovery [27]. In order to address this issue, some studies have proposed discriminative motif discovery, i.e. using negative samples or a background set to increase both the sensitivity and specificity of motif discovery. Some examples of discriminative motif miners are DEME [28] (using a Bayesian framework over alignment columns), MotifHound [29] (hypergeometric test on certain regular expressions in the input data), DLocalMotif [30] (combining motif over-representation, entropy and spatial confinement in motif scoring).

### Motif databases

General-purpose or specialized datasets are dedicated to maintaining a set of experimentally verified motifs from various resources (e.g., gene ontology). ELM [21] as a general-purpose dataset of SLiM, and NLSdb [31] as a specialized database for nuclear-specific motifs is instances of such efforts. Evaluation of mined motifs can be also subjective. Since the extracted motifs do not always exactly match the experimental motifs, residue-level or site-level evaluations have been proposed [26, 32]. Despite great efforts in this area, computational motif mining has remained a challenging task and the state-of-the-art *de novo* approaches have reported relatively low precision and recall scores, even at the residue level [26].

### Protein embedding

Word embedding has been one of the revolutionary concepts in NLP over the recent years and has been shown to be one of the most effective representations in NLP [33, 34, 35]. In particular, skip-gram neural networks combined with negative sampling [36] has resulted in the state-of-the-art performance in a broad range of NLP tasks [35]. Recently, we introduced k-mer-based embedding of biological sequences using skip-gram neural network and negative sampling [5], which became popular for protein feature extraction and has been extended for various classifications of biological sequences [37, 38, 39, 40, 41, 42, 43].

In this work, inspired by unsupervised word segmentation in NLP, we propose a general-purpose segmentation of protein sequences in frequent variable-length sub-sequences called PPE, as a new representation for machine learning tasks. This segmentation is trained once over large protein sequences (Swiss-Prot) and then is applied to a given set of sequences. In this paper, we use this representation for developing a protein motif discovery framework as well as protein sequence embedding.

#### (i) DiMotif

We suggest a discriminative and alignment-free approach for motif discovery that is capable of finding co-occurred motifs. We do not use sequence alignment; instead, we propose the use of general-purpose segmentation of positive and negative input sequences into PPE sequence segments. Subsequently, we use statistical tests to identify the significant discriminative features associated with the positive class, which are our ultimate output motifs. Being alignment-free makes DiMotif, in particular, a favorable choice for the settings where the positive sequences are not necessarily homologous sequences. In the end, we create sets of multi-part motifs using information theoretic measures on the occurrence patterns of motifs on the positive set. We evaluate DiMotif in the detection of the motifs related to 20 types of experimentally verified motifs and also for searching experimentally verified nuclear localization signal (NLS) motifs. The DiMotif achieved a high recall and a reasonable F1 in comparison with the competitive approaches. In addition, we evaluate a shortlist of extracted motifs on the classification of reserved sequences of the same phenotype for integrins, integrin-binding proteins, and biofilm formation proteins, where the phenotype has been detected with a high F1 score. However, a detailed analysis of the motifs and their biophysical properties are beyond the scope of this study, as the main focus is on introducing the method.

#### (ii) ProtVecX

We extend our previously proposed protein vector embedding (Protvec) [5] trained on k-mer segments of the protein sequences to a method of training them on variable-length segments of protein sequences, called ProtVecX. We evaluate our embedding via three protein classification tasks: (i) toxin prediction (binary classification), (ii) subcellular location prediction (four-way classification), and (iii) prediction of enzyme proteins versus non-enzymes (binary classification). We show that concatenation of the raw k-mer distributions with the embedding representations can improve the sequence classification performance over the use of either of k-mers only or embeddings only. In addition, combining of ProtVecX with k-mer occurrence can marginally outperform the use of our originally proposed ProtVec embedding together with k-mer occurrences in toxin and enzyme prediction tasks.

## 2 Material and Methods

### 2.1 Datasets

#### Motif discovery datasets

##### ELM dataset

The eukaryotic linear motif dataset (ELM) is a commonly used resource of experimentally verified SLiMs. The ELM dataset is usually served as the gold standard for the evaluations of *de novo* motif discovery approaches. Due to the long run-time of DLocalMotif (one of the competitive methods) on large datasets, we perform the evaluation on a subset of ELM. In order to cover a variety of settings, we perform the evaluation on 20 motif discovery problems of 5 different motif types: (i) targeting site (TRG), (ii) post-translational modification sites (MOD), (iii) ligand binding sites (LIG), (iv) docking sites (DOC), and (v) degradation sites (DEG). From each category we randomly select 4 sub-types, which are as distinctive as possible (based on the text similarities in their title). Since the ELM dataset only provides a few accession IDs for each motif sub-types, to obtain more data for the domain-specific segmentation, we expand the positive set using NCBI BLAST. Since our method (DiMotif) and DLocalMotif are both instances of discriminative motif discovery methods, we have to create a background/negative as well. For this purpose, we randomly select sequences from UniRef50 ending up a dataset 10 times larger than the whole ELM sequence dataset. Then for each motif type, we calculate the sequence 3-mer representation distance between those randomly selected sequences and the positive set. Subsequently, we sample from the randomly selected sequences based on the cosine distance distribution (considering the same size for the positive and the negative classes). This way the farther sequences from the positive set are more likely to be selected in the background or negative set.

##### Integrin-binding proteins

We extract two positive and negative lists for integrin-binding proteins using the gene ontology (GO) annotation in the UniProt database [44]. For the positive class, we select all proteins annotated with the GO term GO:0005178 (integrin-binding). Removing all redundant sequences results in 2966 protein sequences. We then use 10% of sequences as a reserved set for evaluation and 90% for motif discovery and training purposes. For the negative class, we select a list of proteins sequences which are annotated with the GO term GO:0005515 (protein binding), but they are annotated neither as integrin-binding proteins (GO:0005178) nor the integrin complex (GO:0008305). Since the resulting set are still large, we limit the selection to reviewed Swiss-Prot sequences and filtered redundant sequences, resulting in 20,117 protein sequences, where 25% of these sequences (5029 sequences) are considered as the negative instances for training and validation, and 297 randomly selected instances (equal to 10% of the positive reserved set) as the negative instances for the negative part of the reserved set.

##### Integrin proteins

We extract a list of integrin proteins from the UniProt database by selecting entries annotated with the GO term GO:8305 (integrin complex) that also have integrin as part of their entry name. Removing the redundant sequences results in 112 positive sequences. For the negative sequences, we select sequences which are annotated for transmembrane signaling receptor activity (to be similar to integrins) (GO:4888) but which are neither the integrin complex (GO:8305) nor integrin-binding proteins (GO:0005178). Selection of reviewed Swiss-Prot sequences and removal of redundant proteins results in 1155 negative samples. We use 10% of both the positive and negative sequences as the reserved set for evaluation and 90% for motif discovery and training purposes.

##### Biofilm formation

Similar to integrin-binding proteins, positive and negative lists for biofilm formation are extracted via their GO annotation in UniProt [44]. For the positive class, we select all proteins annotated with the GO term GO:0042710] (biofilm formation). Removing all redundant sequences results in 1450 protein sequences. For the negative class, we select a list of protein sequences annotated within the parent node of biofilm formation in the GO database that is classified as being for a multi-organism cellular process (GO:44764) but not biofilm formation. Since the number of resulting sequences is large, we limit the selection to reviewed Swiss-Prot sequences and filter the redundant sequences, resulting in 1626 protein sequences. Again, we use 10% of both the positive and negative sequences as a reserved set for evaluation and 90% for motif discovery and training purposes.

##### Nuclear localization signals

We use the NLSdb dataset containing nuclear export signals and NLS along with experimentally annotated nuclear and non-nuclear proteins [31]. By using NLSdb annotations from nuclear proteins, we extract a list of proteins experimentally verified to have NLS, ending up with a list of 416 protein sequences. For the negative class, we use the protein sequences in NLSdb annotated as being non-nuclear proteins. NLSdb also contains a list of 3254 experimentally verified motifs, which we use for evaluation purposes.

#### Protein classification datasets

##### Sub-cellular location of eukaryotic proteins

The first dataset we use for protein classification is the TargetP 4-classes dataset of sub-cellular locations. The 4 classes in this dataset are (i) 371 mitochondrial proteins, (ii) 715 pathway or signal peptides, (iii) 1214 nuclear proteins, and (iv) 438 cytosolic protein sequences [45], where the redundant proteins are removed.

##### Toxin prediction

The second dataset we use is the toxin dataset provided by ToxClassifier [46]. The positive set contains 8093 protein sequences annotated in Tox-Prot as being animal toxins and venoms [47]. For the negative class, we choose the ‘Hard’ setting of ToxClassifier [46], where the negative instances are 7043 protein sequences in UniProt which are not annotated in Tox-Prot but are similar to Tox-Prot sequences to some extent.

##### Enzyme detection

On the third we use an enzyme classification dataset. We download two lists of enzyme and non-enzyme proteins (22,168 protein sequences per class) provided by the ‘NEW’ dataset of Deepre [48].

### 2.2 Peptide-pair encoding

#### PPE training

The input to the PPE algorithm is a set of sequences and the output would be segmented sequences and segmentation operations, an ordered list of amino acid merging operations to be applied for segmenting new sequences. At the beginning of the algorithm, we treat each sequence as a list of amino acids. As detailed in Algorithm 1, we then search for the most frequently occurring pair of adjacent amino acids in all input sequences. In the next step, the select pairs of amino acids are replaced by the merged version of the selected pair as a new symbol (a short peptide). This process is continued until we could not find a frequent pattern or we reach a certain vocabulary size (Algorithm 1).

In order to train a general-purpose segmentation of protein sequences, we train the segmentation over the most recent version of the Swiss-Prot database [49], which contained 557,012 protein sequences (step (i) in Figure 2). We continue the merging steps for *T* iterations of Algorithm 1, which ensures that we capture the motifs present with a minimum frequency *f* in all Swiss-Prot sequences (we set the threshold to a minimum of *f* = 10 times, resulting in *T* ≈ 1 million iterations). Subsequently, the merging operations can be applied to any given protein sequences as a general-purpose splitter. Although there exist larger sequence datasets than Swiss-Prot (e.g., UniProt and RefSeq), we decided to use the Swiss-Prot database, due to the high quality, and as computational requirements were less demanding. Principally, PPE segmentations can be generated from any database of interested, given enough time and computational resources.

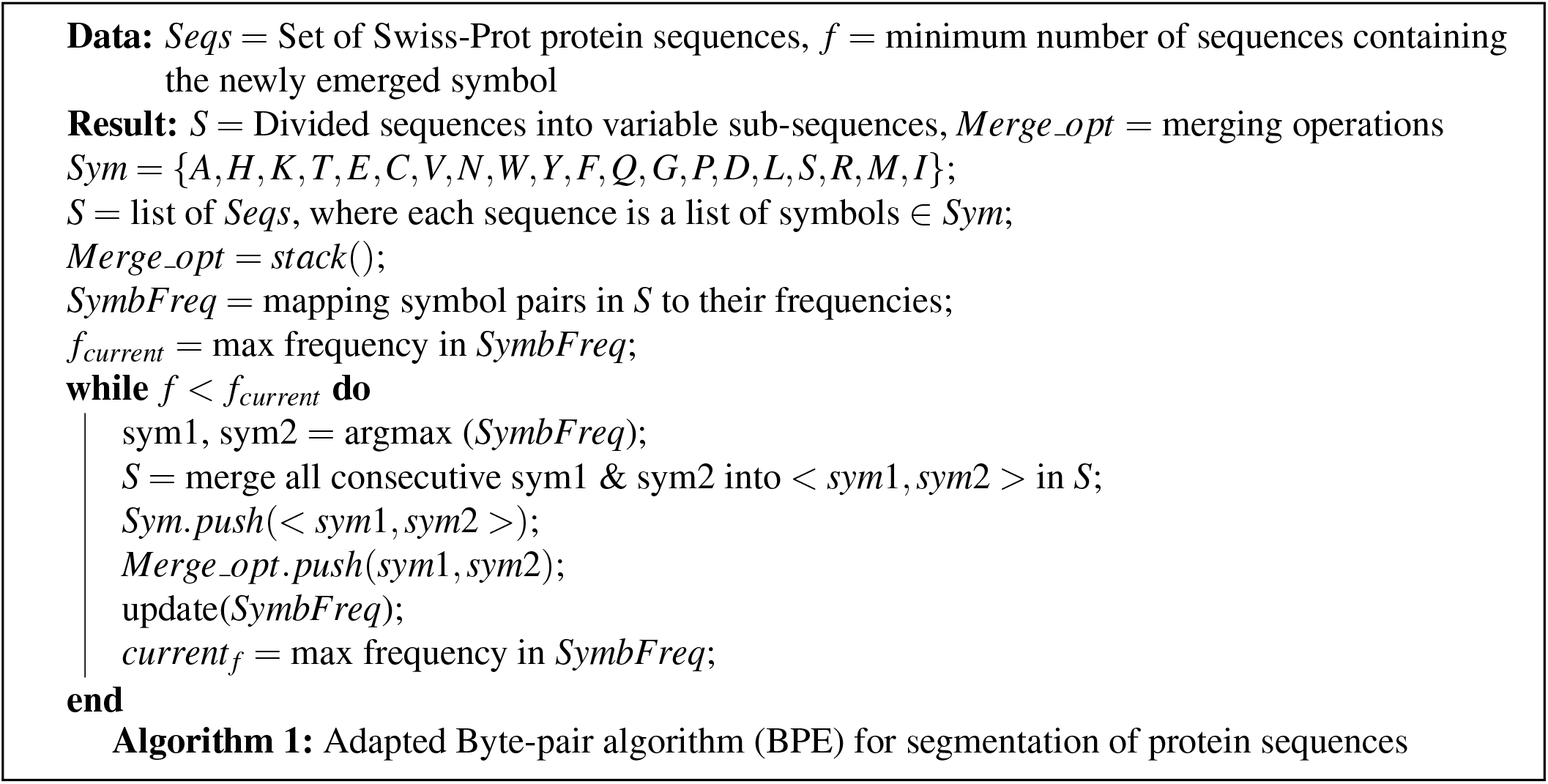

#### Monte Carlo PPE segmentation

The PPE algorithm for a given vocabulary size (which is analogous to the number of merging steps in the training) divides a protein sequence into a unique sequence of sub-sequences. Further merging steps result in enlargement of sub-sequences, which results in having fewer sub-sequences. Such variations can be viewed as multiple valid schemes of sequence segmentation. For certain tasks, it might be useful to consider a protein sequence as a chain of residues and, in some cases, as a chain of large protein domains. Thus, sticking to a single segmentation scheme will result in ignoring important information for the task of interest. In order to address this issue, we propose a sampling framework for estimating the segmentation of a sequence in a probabilistic manner. We sample from the space of possible segmentations for both motif discovery and embedding creation.

Different segmentation schemes for a sequence can be obtained by a varying number of merging steps (*N*) in the PPE algorithm. However, since the algorithm is trained over a large number of sequences, a single merging step will not necessarily affect all sequences, and as we go further with merging steps, fewer sequences are affected by the newly introduced symbol. We estimate the probability density function of possible segmentation schemes with respect to *N* by averaging the segmentation alternatives over 1000 random sequences in Swiss-Prot for *N* ∈ [10000,1000000], with a step size of 10000. For each *N*, we count the average number of introduced symbols relative to the previous step; the average is shown in Figure 1. We use this distribution to draw samples from the vocabulary sizes that affected more sequences (i.e. those introducing more alternative segmentation schemes). To estimate this empirical distribution with a theoretical distribution, we fit a variety of distributions (Gaussian; Laplacian; and *Al pha*, *Beta*, and *Gamma* distributions) using maximum likelihood and found the *Al pha* that fitted the empirical distribution the best. Subsequently, we use the fitted Alpha distribution to draw segmentation samples from a sequence in a Monte Carlo scheme. Consider Φ_*i,j*_ as the *1*1 normalized “bag-of-word” representations in sequence *i*, using the *j^th^* sample from the fitted *Alpha* distribution. Thus, if we have *M* samples, the “bag-of-sub-sequences” representation of the sequence *i* can be estimated as 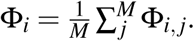.

**Figure 1.**
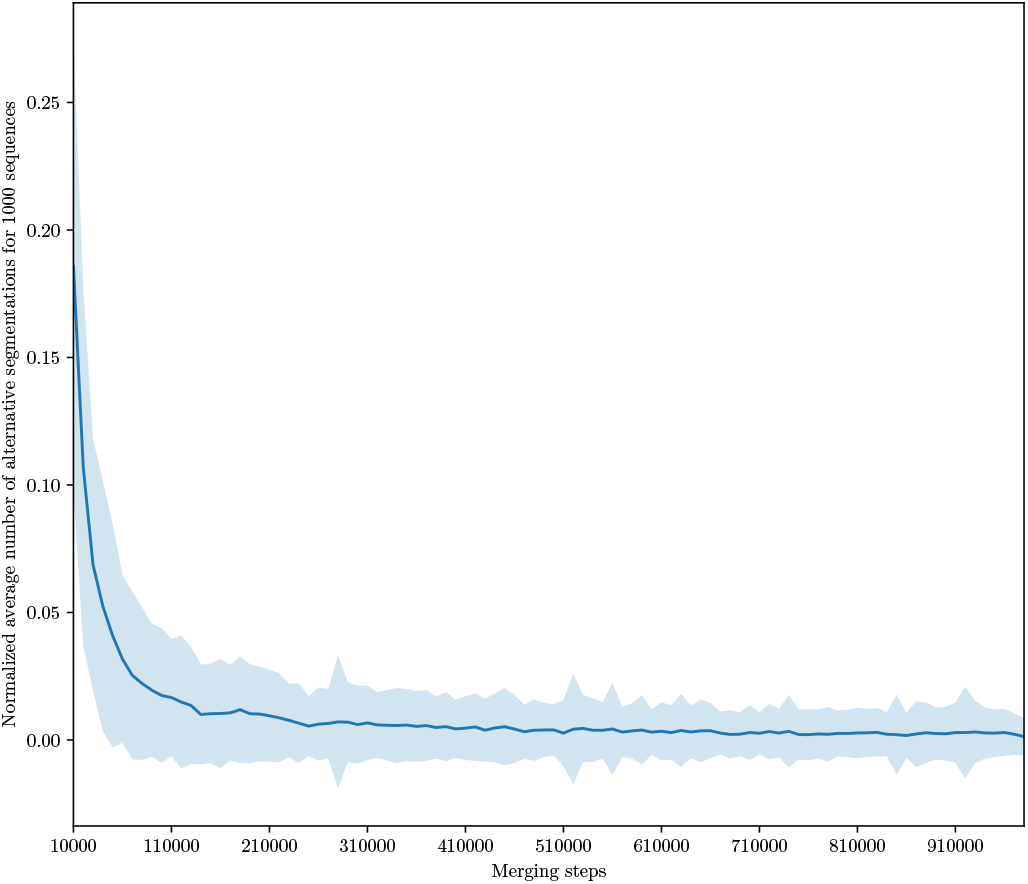
Average number of segmentation alternation per merging steps for 1000 Swiss-prot sequences.

For detecting sequence motifs, we represented each sequence as an average over the count distribution of M samples of segmentation (*M*=100) drawn from the *Alpha* distribution. The alternative is to use only the vocabulary size (e.g., the median of *Alpha*), referred to as the non-probabilistic segmentation in this paper.

### 2.3 DiMotif protein sequence motif discovery

Our proposed method for motif detection, called DiMotif, finds motifs in a discriminative setting over PPE features (step (ii) in Figure 2). We segment all the sequences in the datasets (ignoring their labels or their membership of the train or the test set) with the learned PPE segmentation steps from Swiss-Prot (§2.2) (general-purpose segmentation) or with the learned PPE segmentation steps from a set of positive sequences (domain-specific segmentation). After segmentation, each sequence is represented as bag-of-PPE units. We use a two-sided and FDR corrected *χ*^2^ test to identify significant discriminative features between the positive and the negative (or background) sets. We discard insignificant motifs using a threshold for the p-value of < 0.05. Since we are looking for sequence motifs related to the positive class, we exclude motifs related to the negative class.

**Figure 2.**
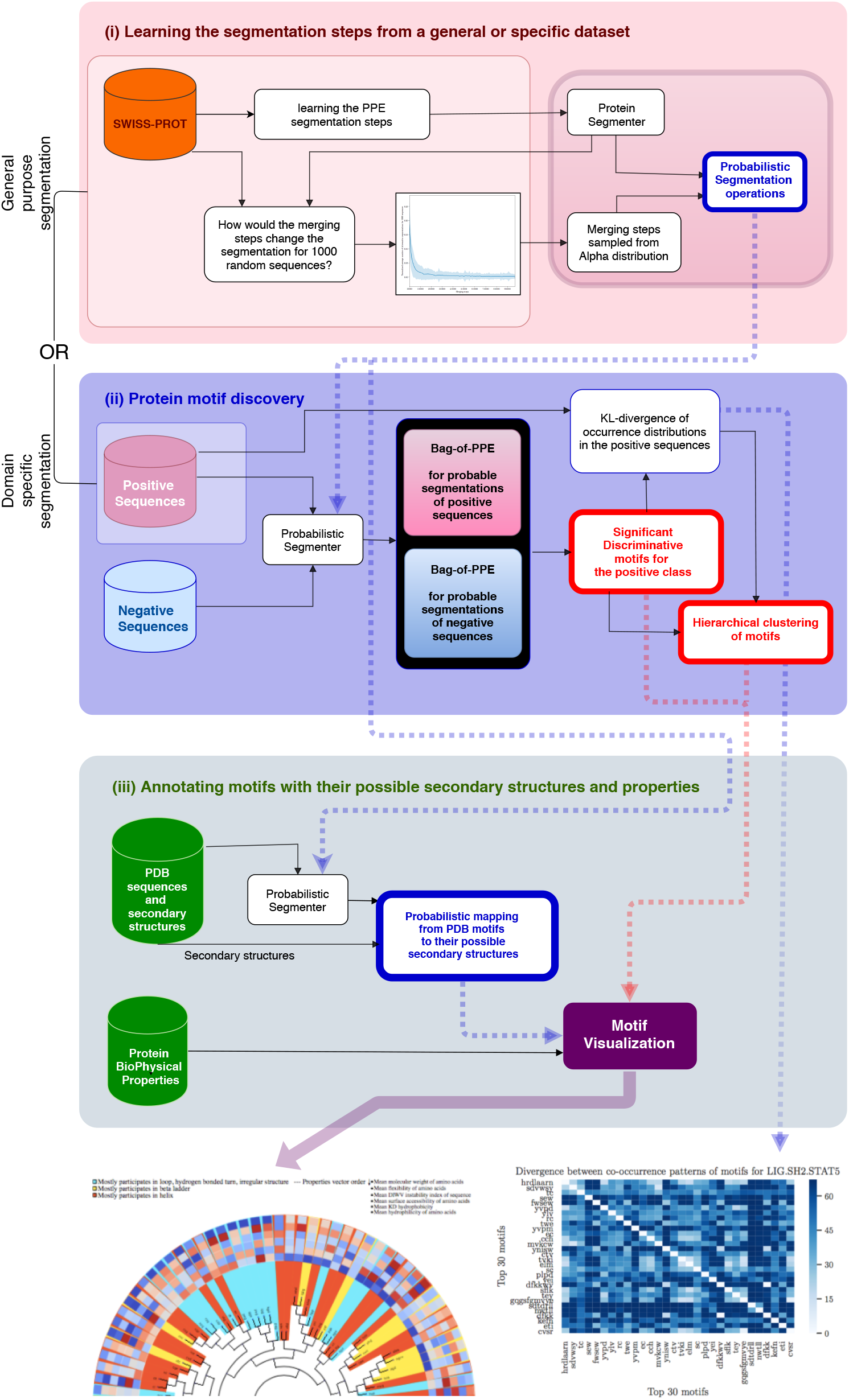
The main steps of DiMotif computational workflow: (i) The PPE segmentation steps can be learned from Swiss-Prot or a domain-specific set of sequences. (ii) These operations are then applied to positive and negative sequences and segment them into smaller sub-sequences. This means that all part of sequences are used till this part. A two-sided and FDR corrected χ^2^ test is then applied to find the sub-sequences (potential motifs) which are significantly related to the positive class, with a threshold p-value < 0.05. We rank the motifs based on their significance and retrieve the top-k (in the evaluation on the ELM dataset *k_m_ax* = 30). (iii) The motifs, their structural and biophysical properties, and their co-occurrence information will be used for visualization.

#### Evaluation on ELM dataset

We compare the DiMotif performance with two recent motif discovery tools: (i) HH-Motif [26] as an instance of non-discriminative methods and (ii) DLocalMotif [30] as an instance of discriminative approaches. We evaluate the performances over the 20 problem settings related to 5 types of motifs in the ELM database. We measure precision and recall of the above methods for detection of the experimentally verified motifs (as true positive). From each method, we use the maximum top 30 retrieved motifs ranked based on their scores. Since finding exact matches are very unlikely and the motifs are only correct to some extent. We report precision and recall for different thresholds on motif sequence matching (50% and 70%). Then we calculate the average precision, recall, and F1 on these different settings. In order to investigate the performance of general-purpose versus domain-specific segmentations in DiMotif, once we used Swiss-Prot segmentation and once we learned the segmentation steps from the set of positive sequences and for both used the probabilistic segmentation schemes.

#### Classification-based evaluation of integrins, integrin-binding proteins, and biofilm formation motifs

In order to evaluate the obtained motifs, we train linear support vector machine classifiers over the training instances but only use motifs related to the positive class among the top 1000 motifs as well as a short list of features. Next, we test the predictive model on a reserved test set. Since the training and testing sets are disjoint, the classification results are indications of motif discovery quality. We use both probabilistic and non-probabilistic segmentation methods to obtain PPE representations of the sequences. We report the precision, recall, and F1 of each classifier’s performance. The average sequence similarities for the top hits between positive samples in the test set and the train set for integrins, integrin binding, and biofilm formation were 35.50±14.41, 40.47±18.15, and 40.13±8.76 respectively. In addition, the average sequence similarities for the best hits for integrins, integrin binding, and biofilm formation were 83.96±11.96, 91.64±11.37, and 71.75±15.79 respectively.

#### NLS motifs search

In the case of NLS motifs, we use the list of 3254 experimentally or manually verified motifs from NLSdb. Thus, in order to evaluate our extracted motifs, we directly compare our motifs with those found in earlier verification. As we cannot evaluate any true positive other than NLSdb this task can be considered as a motif search task. Since for long motifs, finding exact matches is challenging, we report three metrics, the number of motifs with at least three consecutive amino acid overlaps, the number of sequences in the baseline that had a hit with more than 70% overlap (A to B and B to A), and finally the number of exact matches. In addition to Swiss-Prot-based segmentation, in order to see the effect of a specialized segmentation, we also train PPE segmentation over a set of 8421 nuclear protein sequences provided by NLSdb [31] and perform the same evaluation.

#### Kulback-Leibler divergence to find multi-part motifs

As discussed in§ 1, protein motifs can be multi-part patterns, which is ignored by many motif-finding methods. In order to connect the separated parts, we propose to calculate the symmetric Kullback–Leibler (KL) divergence [50] between motifs based on their co-occurrences in the positive sequences as follows:

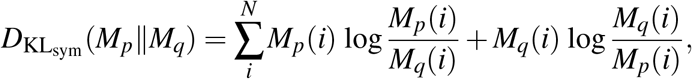

where *M_p_* and *M_q_* are, respectively, the normalized occurrence distributions of motif *p* and motif *q* across all positive samples and *N* is the number of positive sequences. Next, we use the condition of (*D*_KL_sym__ = 0) to find co-occurring motifs splitting the motifs into equivalence classes. Each equivalent class indicates a multi-part or a single-part motif. Since we considered a “bag of motifs” assumption, the parts of multi-part motifs are allowed to be far from each other in the primary sequence.

#### Secondary structure assignment

Using the trained segmentation over the Swiss-Prot sequences, we segment all 385,937 protein sequences in the current version of the PDB [51], where their secondary structure was provided. By segmenting all secondary structures at the same positions as the corresponding sequences, we obtain a mapping from each sequence segment to all its possible secondary structures in the PDB. We use this information in coloring in the visualization of motifs (see the visualizations in §3.1).

##### Motif visualization

For visualization purposes DiMotif clusters motifs based on their co-occurrences in the positive class by using hierarchical clustering over the pairwise symmetric KL divergence. The motifs are then colored based on the most frequent secondary structure they assume in the sequences in the Protein Data Bank (PDB) (step (iii) in Figure 2). For each motif, it visualizes their mean molecular weight, mean flexibility [52], mean instability [53], mean surface accessibility [54], mean kd hydrophobicity [55], and mean hydrophilicity [56] with standardized scores (zero mean and unit variance), where the dark blue is the lowest and the dark red is the highest possible value (see the visualizations in §3.1).

### 2.4 ProtVecX: Extended variable-length protein vector embeddings

We trained the embedding on segmented sequences obtained from Monte Carlo sampling segmentation on the most recent version of the Swiss-Prot database [49], which contains 557,012 protein sequences. Since this embedding is the extended version of ProtVec (a brief introduction to the ProtVec is provided in the Supp. §1), we call it ProtVecX. As explained in §2.2 we segment each sequence with the vocabulary size samples drawn from an *Alpha* distribution. This ensures that we consider multiple ways of segmenting sequences during the embedding training. Subsequently, we train a skip-gram neural network for embedding on the segmented sequences [36]. The skip-gram neural network is analogous to language modeling, which predicts the surroundings (context) for a given textual unit (shown in Figure 3). The skip-gram’s objective is to maximize the following log-likelihood:

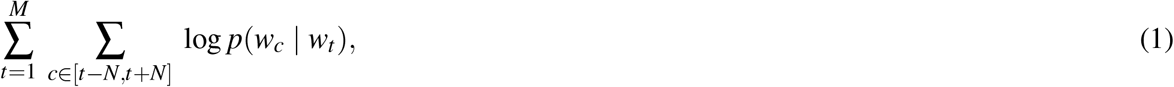

where *N* is the surrounding window size around word *w_t_, c* is the context indices around index *t*, and *M* is the corpus size in terms of the number of available words and context pairs. We parameterize this probability of observing a context word *w_c_* given *w_t_* by using word embedding:

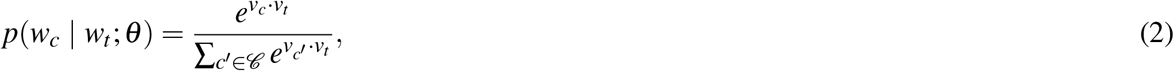

where 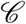 denotes all existing contexts in the training data. However, iterating over all existing contexts is computationally expensive. This issue can be efficiently addressed by using negative sampling. In a negative sampling framework, we can rewrite Equation 1 as follows:

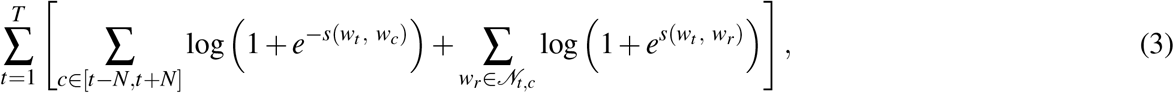

where 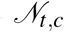 denotes a set of randomly selected negative examples sampled from the vocabulary collection as non-contexts of *w_t_* and *s*(*w_t_, w_c_*) = *v_t_*^T^ · *v_c_* (parameterization with the word vector *v_t_* and the context vector *v_c_*). For training embeddings on PPE units, we used the sub-word level skip-gram, known as fasttext [57]. Fasttext embedding improves the word representations by taking character k-mers of the sub-words into consideration in calculating the embedding of a given word. For instance, if we take the PPE unit *fggagvg* and *k* = 3 as an example, it will be represented by the following character 3-mers and the whole word, where ‘<‘ and ‘>‘ denote the start and the end of a PPE unit:

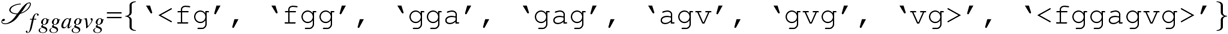

**Figure 3.**
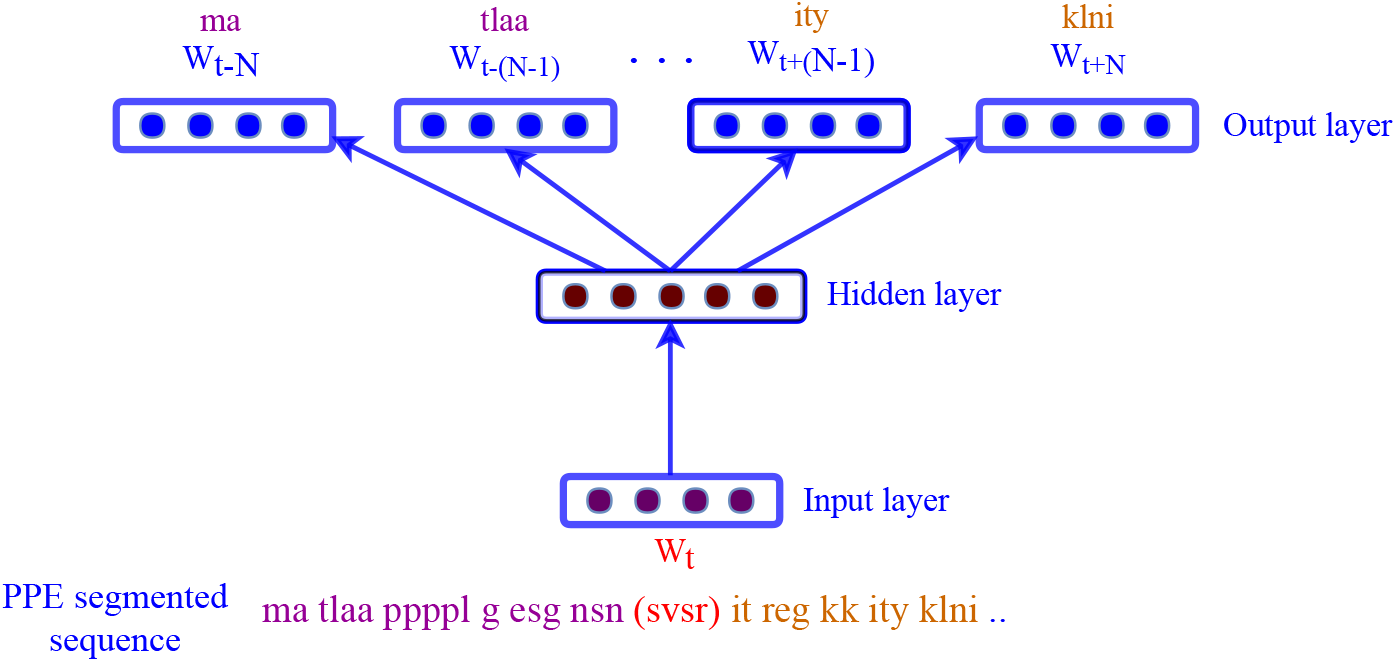
Skip-gram neural network for training language model-based embedding. In this framework the inputs are the segmented sequences and the network is trained to predict the surroundings PPE units.

In the fasttext model, the scoring function will be based on the vector representation of k-mers (2 ≤ *k* ≤ 6) that exist in textual units (PPE units in this case), 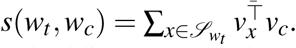.

We used a vector dimension of 500 for the embedding (*v_t_*’s) and a window size of 20 (the vector size and the window size have been selected based on a systematic exploration of parameters in protein classification tasks). A k-mer-based ProtVec of the same vector size and the same window size trained on Swiss-Prot is used for comparison.

#### 2.4.1 Embedding-based classification

For the classification, we use a Multi-Layer-Perceptrons (MLP) neural network architecture with five hidden layers using Rectified Linear Unit (ReLU) as the nonlinear activation function. We use the softmax activation function at the last layer to produce the probability vector that could be regarded as representing posterior probabilities. To avoid overfitting, we perform early stopping and also use dropout at hidden layers. As baseline representations, we use k-mers, ProtVec [5], ProtVecX, and their combinations. For both ProtVec and ProtVecX, the embedding of a sequence is calculated as the summation of its k-mers or PPE unit vectors. We evaluate these representation in three protein classification tasks: (i) toxin prediction (binary classification) with the ‘Hard’ setting in the ToxClassifier database [46], (ii) subcellular location prediction (four-way classification) using the dataset provided by TargetP [45], and (iii) prediction of enzyme proteins versus non-enzymes (binary classification) using the NEW dataset [48]. We report macro-precision, recall, and F-1 score. Macro averaging computes the metrics for each class separately and then simply average over classes. This metric gives equal importance to all categories. In particular, we are interested in macro-F1, which makes a trade-off between precision and recall in addition to treating all classes equally.

## 3 Results

### 3.1 Sequence motifs and evaluation results

#### Detection of experimentally verified motifs in the ELM dataset

A comparison of DiMotif with two existing motif discovery tools, HH-Motif (non-discriminative) and DLocalMotif (discriminative), is provided in Table 1. Since the discovered motifs only partially match the experimentally verified motifs, we measured precision, recall, and F1 scores for two minimum ratios of 50% and 70% sequence matching between the computationally discovered and the experimentally verified motifs (two sets of rows in Table 1 for two minimum sequence matching ratios). DiMotif was used with two different schemes of segmentation: (i) general-purpose segmentation (based-on Swiss-Prot) and (ii) domain-specific segmentation (learned over the sequences in the positive class). Overall, HH-Motif achieved the best F1 scores of 0.39 for 50% sequence matching and F1 of 0.24 for 70% sequence matching. The domain-specific DiMotif obtained F1 of 0.30 for 50% sequence matching and F1 of 0.07 for 70% sequence matching. The general-purpose DiMotif obtained F1 of 0.24 for 50% sequence matching and F1 of 0.05 for 70% sequence matching, while DLocalMotif obtained F1 of 0.16 for 50% sequence matching and F1 of 0.04 for 70% sequence matching. The domain-specific DiMotif achieved the maximum recall of 0.80 for 50% sequence matching and F1 of 0.32 for 70% sequence matching. Having high recall suggests that the DiMotif can be used for short-list creation for further experimental investigations on motifs.

**Table 1.** Comparison of DiMotif, HH-Motif, and DLocalMotif (four sets of columns) performances in detection of experimentally verified motifs in the ELM dataset. Two versions of DiMotif were used: (i) using Swiss-Prot segmentation and (ii) using a domain specific segmentation. The performances are reported for 50% and 70% ratios of sequence matching (two sets of rows) between the verified and the discovered motifs.

|  |  | HH-MOTIF |  |  | DiMotif (Domain-specific) |  |  | DiMotif (SWISS-Prot-based) |  |  | DLocalMotif |  |  |
| --- | --- | --- | --- | --- | --- | --- | --- | --- | --- | --- | --- | --- | --- |
| Motif discovery setting |  | Precision | Recall | F1 score | Precision | Recall | F1 score | Precision | Recall | F1 score | Precision | Recall | F1 score |
| Matching condition: 50% overlap between motifs | DEG APCC KENBOX 2 | 0.80 | 1.00 | 0.89 | 0.07 | 0.94 | 0.13 | 0.26 | 1.00 | 0.41 | 0.23 | 0.81 | 0.36 |
|  | DEG CRL4 CDT2 1 | 0.60 | 1.00 | 0.75 | 0.23 | 0.83 | 0.36 | 0.00 | 0.00 | 0.00 | 0.00 | 0.00 | 0.00 |
|  | DEG Kelch KLHL3 1 | 0.00 | 0.00 | 0.00 | 0.03 | 1.00 | 0.06 | 0.00 | 0.00 | 0.00 | 0.00 | 0.00 | 0.00 |
|  | DEG SPOP SBC 1 | 0.00 | 0.00 | 0.00 | 0.03 | 0.14 | 0.05 | 0.20 | 0.71 | 0.31 | 0.07 | 0.86 | 0.13 |
|  | DOC AGCK PIF 3 | 0.00 | 0.00 | 0.00 | 0.00 | 0.00 | 0.00 | 0.00 | 0.00 | 0.00 | 0.00 | 0.00 | 0.00 |
|  | DOC MAPK 2 | 0.59 | 1.00 | 0.74 | 0.17 | 1.00 | 0.29 | 0.33 | 1.00 | 0.50 | 0.00 | 0.00 | 0.00 |
|  | DOC PIKK 1 | 0.33 | 0.50 | 0.40 | 0.17 | 1.00 | 0.29 | 0.10 | 0.75 | 0.18 | 0.00 | 0.00 | 0.00 |
|  | DOC USP7 MATH 1 | 0.00 | 0.00 | 0.00 | 0.30 | 0.60 | 0.40 | 0.20 | 0.60 | 0.30 | 0.00 | 0.00 | 0.00 |
|  | LIG 14-3-3 2 | 0.08 | 0.28 | 0.13 | 0.30 | 1.00 | 0.46 | 0.17 | 0.43 | 0.24 | 0.10 | 0.43 | 0.16 |
|  | LIG Mtr4 Air2 1 | 0.00 | 0.00 | 0.00 | 0.13 | 1.00 | 0.23 | 0.00 | 0.00 | 0.00 | 0.00 | 0.00 | 0.00 |
|  | LIG SH2 STAT5 | 0.29 | 0.38 | 0.33 | 0.47 | 1.00 | 0.64 | 0.30 | 1.00 | 0.46 | 0.00 | 0.00 | 0.00 |
|  | LIG TYR ITIM | 0.25 | 0.17 | 0.20 | 0.27 | 0.83 | 0.41 | 0.17 | 0.50 | 0.25 | 0.27 | 0.67 | 0.38 |
|  | MOD CDK 1 | 0.30 | 0.70 | 0.42 | 0.20 | 0.90 | 0.33 | 0.30 | 0.90 | 0.45 | 0.53 | 0.80 | 0.64 |
|  | MOD NEK2 1 | 0.33 | 0.66 | 0.44 | 0.17 | 1.00 | 0.29 | 0.04 | 0.33 | 0.07 | 0.00 | 0.00 | 0.00 |
|  | MOD SPalmitoyl 4 | 1.00 | 1.00 | 1.00 | 0.17 | 1.00 | 0.29 | 0.14 | 1.00 | 0.25 | 0.53 | 1.00 | 0.69 |
|  | MOD SUMO rev 2 | 0.17 | 0.05 | 0.08 | 0.37 | 0.79 | 0.50 | 0.37 | 0.53 | 0.44 | 0.30 | 0.21 | 0.25 |
|  | TRG AP2beta CARGO 1 | 0.50 | 1.00 | 0.67 | 0.10 | 0.50 | 0.17 | 0.05 | 0.75 | 0.09 | 0.03 | 0.25 | 0.05 |
|  | TRG ER KDEL 1 | 0.60 | 1.00 | 0.75 | 0.37 | 1.00 | 0.54 | 0.43 | 1.00 | 0.60 | 0.23 | 0.20 | 0.21 |
|  | TRG LysEnd APsAcLL 3 | 1.00 | 1.00 | 1.00 | 0.20 | 1.00 | 0.33 | 0.17 | 1.00 | 0.29 | 0.00 | 0.00 | 0.00 |
|  | TRG NES CRM1 1 | 0.08 | 0.06 | 0.07 | 0.17 | 0.56 | 0.26 | 0.00 | 0.00 | 0.00 | 0.00 | 0.00 | 0.00 |
| Macro-Averaged Metrics |  | 0.35 | 0.49 | 0.39 | 0.20 | 0.80 | 0.30 | 0.16 | 0.57 | 0.24 | 0.11 | 0.26 | 0.16 |
| Matching condition: 70% overlap between motifs | DEG APCC KENBOX 2 | 0.52 | 0.88 | 0.65 | 0.03 | 0.94 | 0.06 | 0.10 | 0.38 | 0.16 | 0.03 | 0.06 | 0.04 |
|  | DEG CRL4 CDT2 1 | 0.52 | 1.00 | 0.68 | 0.00 | 0.00 | 0.00 | 0.00 | 0.00 | 0.00 | 0.00 | 0.00 | 0.00 |
|  | DEG Kelch KLHL3 1 | 0.00 | 0.00 | 0.00 | 0.03 | 0.50 | 0.06 | 0.00 | 0.00 | 0.00 | 0.00 | 0.00 | 0.00 |
|  | DEG SPOP SBC 1 | 0.00 | 0.00 | 0.00 | 0.00 | 0.00 | 0.00 | 0.17 | 0.71 | 0.27 | 0.00 | 0.00 | 0.00 |
|  | DOC AGCK PIF 3 | 0.00 | 0.00 | 0.00 | 0.00 | 0.00 | 0.00 | 0.00 | 0.00 | 0.00 | 0.00 | 0.00 | 0.00 |
|  | DOC MAPK 2 | 0.24 | 1.00 | 0.39 | 0.07 | 1.00 | 0.13 | 0.07 | 0.00 | 0.00 | 0.00 | 0.00 | 0.00 |
|  | DOC PIKK 1 | 0.00 | 0.00 | 0.00 | 0.00 | 0.00 | 0.00 | 0.00 | 0.00 | 0.00 | 0.00 | 0.00 | 0.00 |
|  | DOC USP7 MATH 1 | 0.00 | 0.00 | 0.00 | 0.03 | 0.10 | 0.05 | 0.13 | 0.30 | 0.18 | 0.00 | 0.00 | 0.00 |
|  | LIG 14-3-3 2 | 0.00 | 0.00 | 0.00 | 0.00 | 0.00 | 0.00 | 0.00 | 0.00 | 0.00 | 0.00 | 0.00 | 0.00 |
|  | LIG Mtr4 Air2 1 | 0.00 | 0.00 | 0.00 | 0.07 | 1.00 | 0.13 | 0.00 | 0.00 | 0.00 | 0.00 | 0.00 | 0.00 |
|  | LIG SH2 STAT5 | 0.00 | 0.00 | 0.00 | 0.13 | 0.50 | 0.21 | 0.13 | 0.25 | 0.17 | 0.00 | 0.00 | 0.00 |
|  | LIG TYR ITIM | 0.00 | 0.00 | 0.00 | 0.03 | 0.16 | 0.05 | 0.00 | 0.00 | 0.00 | 0.13 | 0.33 | 0.19 |
|  | MOD CDK 1 | 0.30 | 0.20 | 0.24 | 0.03 | 0.30 | 0.05 | 0.00 | 0.00 | 0.00 | 0.07 | 0.20 | 0.10 |
|  | MOD NEK2 1 | 0.00 | 0.00 | 0.00 | 0.03 | 0.33 | 0.05 | 0.00 | 0.00 | 0.00 | 0.00 | 0.00 | 0.00 |
|  | MOD SPalmitoyl 4 | 1.00 | 1.00 | 1.00 | 0.30 | 0.20 | 0.24 | 0.00 | 0.00 | 0.00 | 0.13 | 0.40 | 0.20 |
|  | MOD SUMO rev 2 | 0.00 | 0.00 | 0.00 | 0.10 | 0.16 | 0.12 | 0.03 | 0.05 | 0.04 | 0.00 | 0.00 | 0.00 |
|  | TRG AP2beta CARGO 1 | 0.00 | 0.00 | 0.00 | 0.00 | 0.00 | 0.00 | 0.00 | 0.00 | 0.00 | 0.00 | 0.00 | 0.00 |
|  | TRG ER KDEL 1 | 0.60 | 1.00 | 0.75 | 0.13 | 0.80 | 0.22 | 0.13 | 0.80 | 0.22 | 0.23 | 0.20 | 0.21 |
|  | TRG LysEnd APsAcLL 3 | 1.00 | 1.00 | 1.00 | 0.03 | 0.33 | 0.05 | 0.00 | 0.00 | 0.00 | 0.00 | 0.00 | 0.00 |
|  | TRG NES CRM1 1 | 0.00 | 0.00 | 0.00 | 0.00 | 0.00 | 0.00 | 0.00 | 0.00 | 0.00 | 0.00 | 0.00 | 0.00 |
| Macro-Averaged Metrics |  | 0.21 | 0.30 | 0.24 | 0.05 | 0.32 | 0.07 | 0.04 | 0.12 | 0.05 | 0.03 | 0.06 | 0.04 |

#### Classification-based evaluation of integrins, integrin-binding, and biofilm formation motifs

The performances of machine learning classifiers in phenotype prediction using the extracted motifs as features are provided in Table 2 evaluated in both 10-fold cross-validation scheme, as well as in classifying unseen reserved sequences. Both probabilistic and non-probabilistic segmentation methods have been used to obtain PPE motifs. However, from the top extracted motifs only motifs associated with the positive class are used as features (representation column). For each classification setting, we report precision, recall, and F1 scores. The trained classifiers over the extracted motifs associated with the positive class could reliably predict the reserved integrins, integrin-binding proteins, and biofilm formation proteins with F1 scores of 0.89, 0.89, and 0.75 respectively. As described in §2.1 the sequences with certain degrees of redundancy were already removed and the training data and the reserved sets do not overlap. Thus, being able to predict the phenotype over the reserved sets with high F1 scores shows the quality of motifs extracted by DiMotif. This confirms that the extracted motifs are specific and inclusive enough to detect the phenotype of interest among an unseen set of sequences.

**Table 2.** Evaluation of protein sequence motifs mined via PPE motif discovery for classification of integrin-binding proteins and biofilm formation-associated proteins. Support Vector Machine classifiers are tuned and evaluated in a stratified 10-fold cross-validation setting and then tested on a separate reserved dataset.

| Dataset | Probabilistic Segmentation | Representation | 10-fold cross-validation |  |  | Performance on the test set |  |  |
| --- | --- | --- | --- | --- | --- | --- | --- | --- |
|  |  |  | Precision | Recall | F1 | Precision | Recall | F1 |
| Integrin-Binding | True | top 1000 (998 positive) | 0.88 | 0.85 | 0.87 | 0.91 | 0.85 | 0.88 |
|  |  | top-100 (100 positive) | 0.73 | 0.66 | 0.69 | 0.84 | 0.67 | 0.75 |
|  | False | top 1000 (982 positive) | 0.91 | 0.87 | 0.89 | 0.93 | 0.86 | **0.89** |
|  |  | top-100 (100 positive) | 0.73 | 0.68 | 0.70 | 0.84 | 0.67 | 0.75 |
| Integrins | True | top 1000 (1000 positive) | 0.94 | 0.76 | 0.84 | 1 | 0.75 | 0.86 |
|  |  | top-100 (100 positive) | 0.91 | 0.82 | 0.86 | 0.83 | 0.83 | **0.83** |
|  | False | top 1000 (996 positive) | 0.96 | 0.82 | 0.89 | 1 | 0.83 | 0.91 |
|  |  | top-100 (100 positive) | 0.88 | 0.83 | 0.86 | 0.9 | 0.75 | 0.82 |
| Biofilm formation | True | top-1000 (103 positive) | 0.89 | 0.67 | 0.76 | 0.82 | 0.56 | 0.72 |
|  |  | top-500 (48 positive) | 0.81 | 0.71 | 0.76 | 0.76 | 0.71 | **0.73** |
|  | False | top 1000 (53 positive) | 0.79 | 0.67 | 0.73 | 0.74 | 0.66 | 0.70 |
|  |  | top-500 (26 positive) | 0.78 | 0.67 | 0.72 | 0.73 | 0.65 | 0.69 |

For integrin and biofilm formation, the probabilistic segmentation helps in predictions of the reserved dataset. This suggests that multiple views of segmenting sequences allows the statistical feature selection model to be more inclusive in observing possible motifs. Picking a smaller fraction of positive class motifs still resulted in a high F1 for the test sets. For biofilm formation, the probabilistic segmentation improved the classification F1 score from 0.72 to 0.73 when only 48 motifs were used, where single segmentation even using more features obtained an F1 score of 0.70 (Table 2). This classification result suggests that the only 48 motifs mined from the training set are enough to detect bioform formation proteins in the test set. Thus, such a combination can be a good representative of biofilm formation motifs.

#### Literature-based evaluation of NLS motifs

Since NLSdb provided us with an extensive list of experimentally verified NLS motifs, we evaluated the extracted motifs by measuring their overlap with NLSdb instead of using a classification-based evaluation. However, as discussed in §1 such a comparison can be very challenging. One reason is that different methods and technologies vary in their resolutions in specifying the motif boundaries. In addition, the motifs extracted by the computational methods may also contain some degrees of false negatives and false positives. Thus, instead of reporting exact matches in the experimentally verified set, we report how many of 3254 motifs in NLSdb are verified by our approach using three different degrees of similarity (medium overlap, large overlap, and exact match). The performance of DiMotif for both probabilistic segmentation and non-probabilistic segmentation are provided in Table 3. In order to investigate the performance of phenotype-specific versus general purpose segmentation, we also report the results based on segmentation that is learned from nuclear proteins, in addition to Swiss-Prot based segmentation (which is supposed to general purpose). Training the segmentation on nuclear proteins resulted in slightly better, but still competitive to the general-purpose Swiss-Prot segmentation. This result shows that the segmentation steps learned from Swiss-Prot can be considered as a general segmentation, which is important for low resource settings, i.e. the problem setting that the number of positive samples is relatively small. Similar to integrins and biofilm formation related proteins, the probabilistic segmentation has been more successful in detecting experimentally verified NLS motifs as well (Table 3).

**Table 3.** Evaluation of the significant nuclear localization signal (NLS) patterns against 3254 experimentally identified motifs. The results are provided for both general purpose and domain-specific segmentation of sequences.

| PPE training dataset | Probabilistic Segmentation | Medium overlap: Overlapping hits (> 3) | Large overlap: > 70% sequence overlap | Number of exact matches |
| --- | --- | --- | --- | --- |
| Swiss-Prot (General purpose) | True | 3253 | 337 | 37 |
| Swiss-Prot (General purpose) | False | 3162 | 107 | 15 |
| Nuclear (Domain-specific) | True | 3253 | 381 | 42 |
| Nuclear (Domain-specific) | False | 3198 | 137 | 21 |

#### DiMotif Visualization

The top extracted motifs are visualized using DiMotif software and are provided for interested readers, related to integrin-binding proteins (Figure 4), biofilm formation (Figure 5), and integrin complexes (Figure 6). In these visualizations, motifs are clustered according to their cooccurrences within the positive set, i.e. if two motifs tend to occur together (not necessarily close in the linear chain) in these hierarchical clustering they are in a close proximity. In addition, each motif is colored based on the most frequent secondary structure that this motif can assume in all existing PDB structures (described in §2.3), the blue background shows loop, hydrogen bound or irregular structures, the yellow background shows beta ladders, and red background shows helical structures. Furthermore, to facilitate the interpretation of the found motifs, DiMotif provides a heatmap representation of biophysical properties related to each motif, namely molecular weight, flexibility, instability, surface accessibility, kd hydrophobicity, and mean hydrophilicity with standardized scores (zero mean and unit variance) the dark blue is the lowest and the dark red is the highest possible value. Normalized scores allow for an easier visual comparison. For instance, interestingly in most cases in the trees (Figure 4, Figure 5, and Figure 6), the neighbor motifs (co-occurring motifs) agree in their frequent secondary structures. Furthermore, some neighbor motifs agree in some provided biophysical properties. Such information can assist biologists and biophysicists to make hypotheses about the underlying motifs and mechanisms for further experiments. A detailed serious biophysical investigation of the extracted motifs is beyond the scope of this study. However, as an example, for integrin-binding proteins, the RGD motif, the most well-known integrin-binding motif was among the most significant motifs in our approach [58, 59, 60]. Other known integrin-binding motifs were also among the most significant ones, such as RLD [59], KGD (the binding site for the *αIIβ*3 integrins [61]), GPR (the binding site for *α_x_β*2 [60]), LDT (the binding site for *α_4_β*7 [60]), QIDS (the binding site for *α_4_β* 1 [60]), DLLEL (the binding site for *α_v_β*6 [60]), [tldv,rldvv,gldvs] (similar motifs to LDV, the binding site for *α_4_β* 1 [58]), rgds [62], as well as the PEG motif [63].

**Figure 4.**
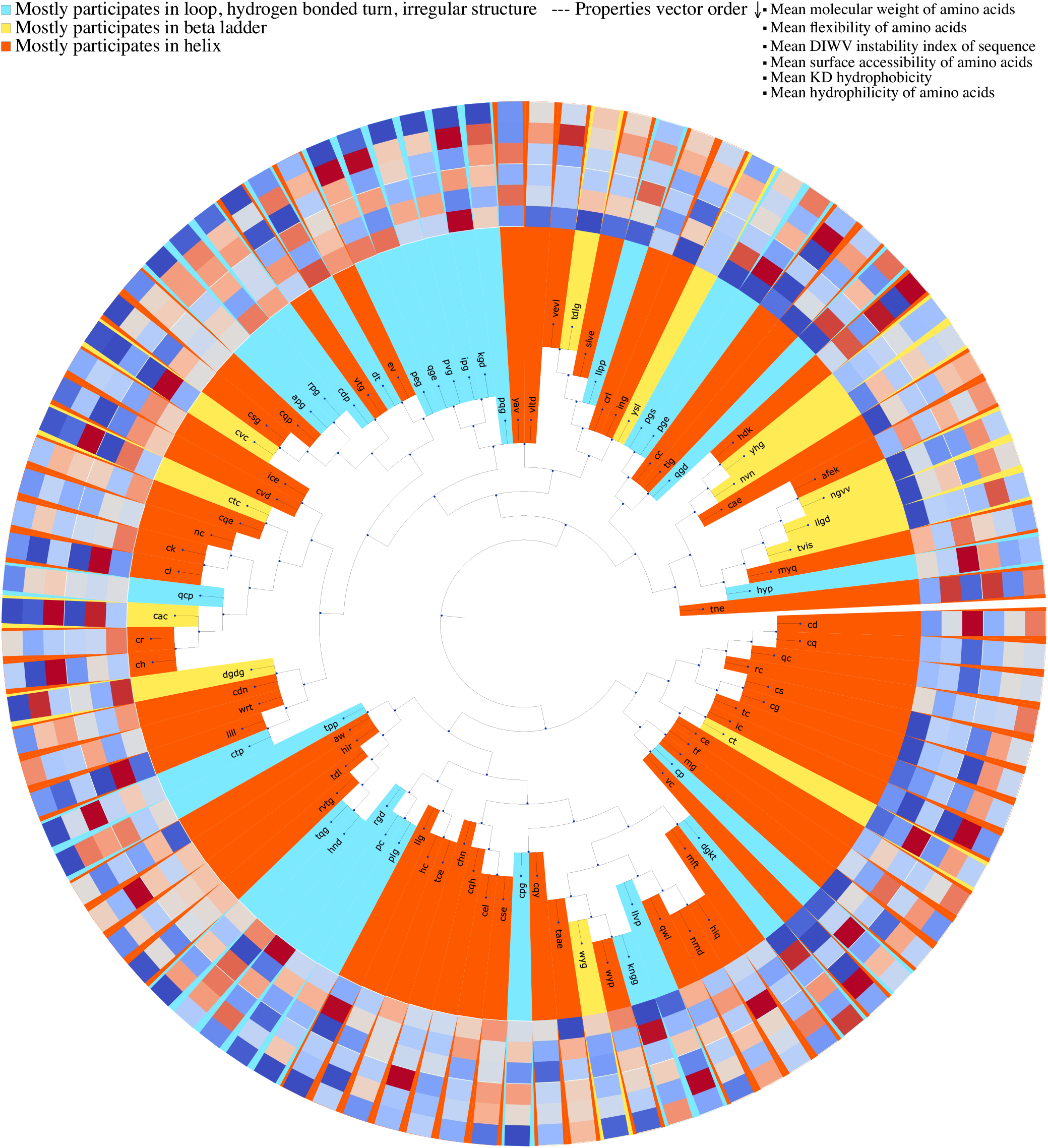
Clustering of integrin-binding-specific motifs based on their occurrence in the annotation proteins. Each motif is colored based on the most frequent secondary structure it assumes in the Protein Data Bank (PDB) structure out of all PDB sequences. For each motif the biophysical properties are provided in a heatmap visualization, which shows from outer ring to inner ring: the mean molecular weight, mean flexibility, mean instability, mean surface accessibility, mean kd hydrophobicity, and mean hydrophilicity with standardized scores (zero mean and unit variance), where the dark blue is the lowest and the dark red is the highest possible value. Motifs are clustered based on their co-occurrences in the integrin-binding proteins.

**Figure 5.**
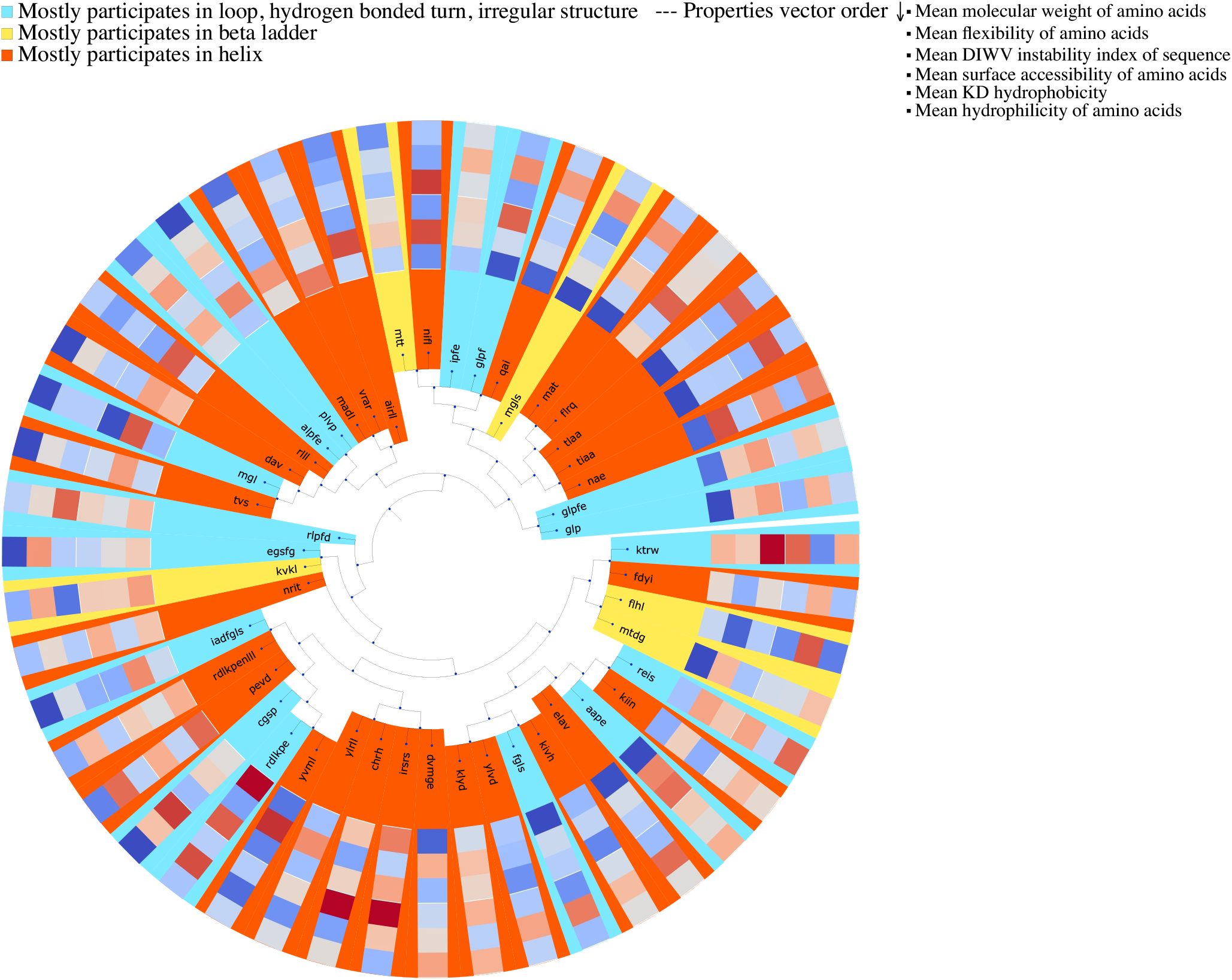
Clustering of biofilm formation-specific motifs based on their occurrence in the annotation proteins. Each motif is colored based on the most frequent secondary structure it assumes in the Protein Data Bank (PDB) structure out of all PDB sequences. For each motif the biophysical properties are provided in a heatmap visualization, which shows from outer ring to inner ring: the mean molecular weight, mean flexibility, mean instability, mean surface accessibility, mean kd hydrophobicity, and mean hydrophilicity with standardized scores (zero mean and unit variance), where the dark blue is the lowest and the dark red is the highest possible value. Motifs are clustered based on their co-occurrences in the biofilm formation proteins.

**Figure 6.**
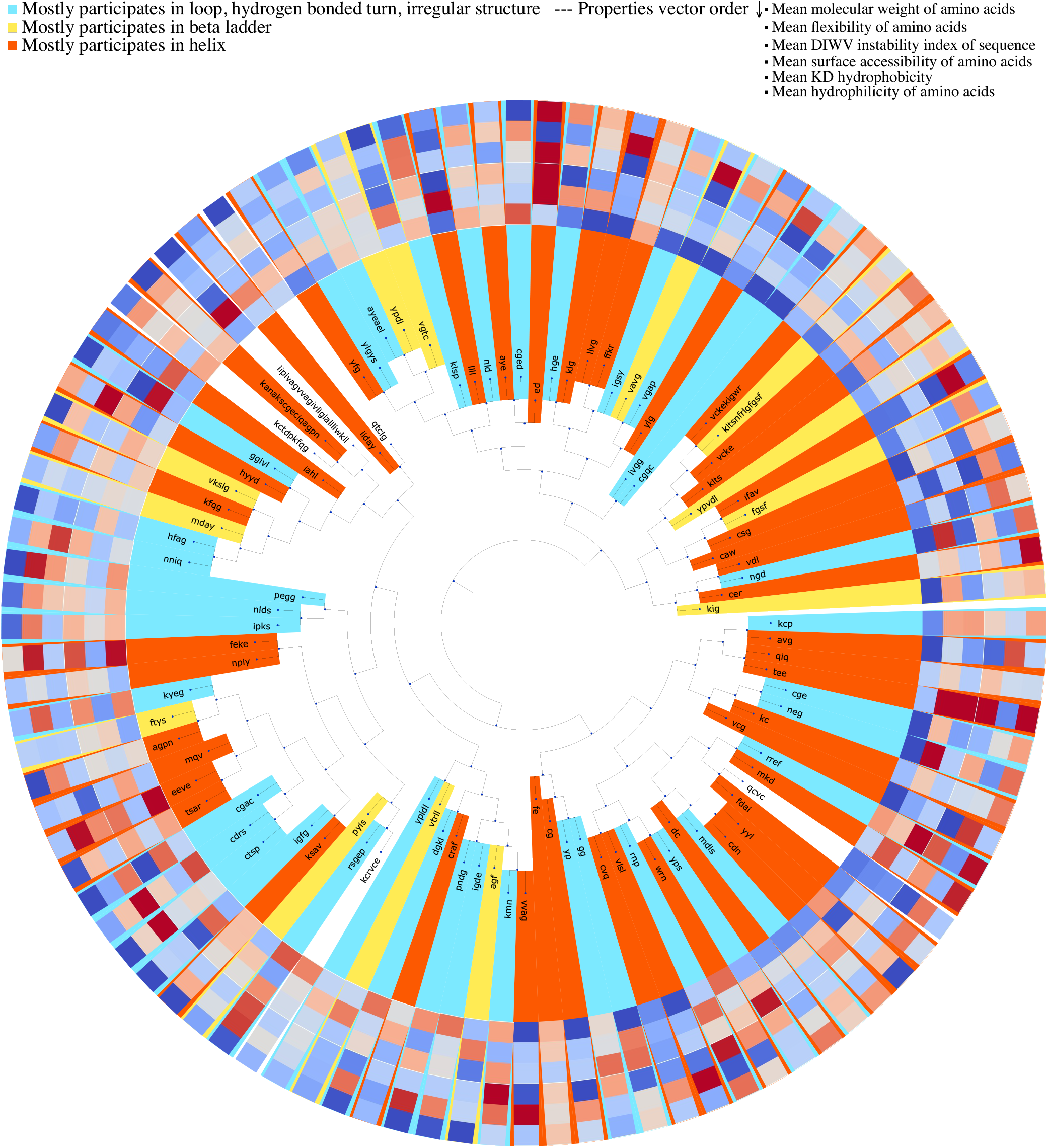
Clustering of integrin-related motifs based on their occurrence in the annotation proteins. Each motif is colored based on the most frequent secondary structure it assumes in the Protein Data Bank (PDB) structure out of all PDB sequences. For each motif the biophysical properties are provided in a heatmap visualization, which shows from outer ring to inner ring: the mean molecular weight, mean flexibility, mean instability, mean surface accessibility, mean kd hydrophobicity, and mean hydrophilicity with standardized scores (zero mean and unit variance), where the dark blue is the lowest and the dark red is the highest possible value. Motifs are clustered based on their co-occurrences in the integrin proteins.

### 3.2 Results of protein classification tasks using embedding

Protein classification results for venom toxins, subcellular location, and enzyme predictions using deep MLP neural network on top of different combinations of features are provided in Table 4. In all these three tasks, combining the embeddings with raw k-mer distributions improves the classification performances (Table 4). This result suggests that k-mers can be more specific than embeddings for protein classification. However, embeddings can provide complementary information to the k-mers and improve the classification performances. Combining 3-mers with either ProtVecX or ProtVec embedding performed very competitively; even for sub-cellular prediction tasks, ProtVec performs slightly better. However, combining 3-mers with ProtVecX resulted in higher F1 scores for enzyme classification and toxin protein prediction. In our previously proposed ProtVec paper [5] as well as other embedding-based protein classification papers [43], embeddings have been used as the only representation. However, the presented results in Table 4 suggest that k-mer representation, although is a simple approach but is a tough-to-beat baseline in classification tasks. The ProtVec and ProtVecX embeddings only have added value when they are combined with the raw k-mer representations.

**Table 4.** Comparing k-mers, ProtVec, and ProtVecX and their combinations in protein classification tasks. Deep MLP neural network has been used as the classifier.

| Dataset | Representation | 5 fold cross-validation |  |  |
| --- | --- | --- | --- | --- |
|  |  | macro-Precision | macro-Recall | macro-F1 |
| Venom toxin prediction | 3-mer | 0.89 | 0.89 | 0.89 |
|  | ProtVec | 0.88 | 0.88 | 0.88 |
|  | ProtVecX | 0.88 | 0.88 | 0.88 |
|  | 3-mer + ProtVec | 0.90 | 0.89 | 0.89 |
|  | 3-mer + ProtVecX | 0.90 | 0.90 | **0.90** |
| Subcellular location prediction | 3-mer | 0.65 | 0.59 | 0.60 |
|  | ProtVec | 0.60 | 0.57 | 0.58 |
|  | ProtVecX | 0.57 | 0.57 | 0.57 |
|  | 3-mer + ProtVec | 0.68 | 0.60 | **0.62** |
|  | 3-mer + ProtVecX | 0.66 | 0.60 | 0.61 |
| Enzyme prediction | 3-mer | 0.70 | 0.73 | 0.71 |
|  | ProtVec | 0.68 | 0.70 | 0.69 |
|  | ProtVecX | 0.69 | 0.71 | 0.70 |
|  | 3-mer + ProtVec | 0.70 | 0.73 | 0.71 |
|  | 3-mer + ProtVecX | 0.71 | 0.73 | **0.72** |

## 4 Conclusions

We proposed a new unsupervised method of feature extraction from protein sequences. Instead of fixed-length k-mers, we segmented sequences into the most frequent variable-length sub-sequences, inspired by BPE, a data compression algorithm. These sub-sequences were then used as features for downstream machine learning tasks. As a modification to the original BPE algorithm, we defined a probabilistic segmentation by sampling from the space of possible vocabulary sizes. This allows for considering multiple ways of segmenting a sequence into sub-sequences. The main purpose of this work was to introduce a variable-length segmentation of sequences, similar to word tokenization in natural languages. In particular, we introduced (i) DiMotif as an alignment-free discriminative protein sequence motif miner, as well as (ii) ProtVecX, a variable-length extension of protein sequence embedding.

We compared DiMotif against two recent existing tools for motif discovery: HH-Motif as an instance of non-discrminative methods, and DLocalMotif as an instance of discriminative methods. We compared the performances in the detection of 20 distinct sub-types of experimentally verified motifs. HH-Motif which uses HMM over multiple sequence alignment achieved the best average F1 and the DiMotif with domain-specific segmentation achieved the second best F1. DiMotif achieved the highest recall, making it an ideal tool for finding a list of candidates for further experimental verifications. Furthermore, We evaluated DiMotif by extracting motifs related to (i) integrins, (ii) integrin-binding proteins, and (iii) biofilm formation. We showed that the extracted motifs could reliably detect reserved sequences of the same phenotypes, as indicated by their high F1 scores. We also showed that DiMotif could reasonably detect experimentally identified motifs related to nuclear localization signals. By using KL divergence between the distribution of motifs in the positive sequences, DiMotif is capable of outputting multi-part motifs. A detailed biophysical interpretation of the motifs is beyond the scope of this work. However, the tree visualization of DiMotif as a tool can help biologists to come up with hypotheses about the motifs for further experiments. In addition, although homologous sequences in Swiss-Prot have indirectly contributed in DiMotif segmentation scheme, unlike conventional motif discovery algorithms, DiMotif does not directly use multiple sequence alignment information. Thus, it can be widely used in cases motifs need to be found from a set of non-homologous sequences.

We proposed ProtVecX embedding trained on sub-sequences in the Swiss-Prot database. We demonstrated that combining the raw k-mer distributions with the embedding representations can improve the sequence classification performance compared with using either k-mers only or embeddings only. In addition, combining ProtVecX with k-mer occurrences outperformed ProtVec embedding combined with k-mer occurrences for toxin and enzyme prediction tasks. Our results suggest that the recent works in the literature including our previously proposed ProtVec missed serving k-mer representation as a baseline, which is a tough-to-beat baseline. We show that embedding can be used as complementary information to the raw k-mer distribution and their added value is expressed when they are combined with k-mer features.

In this paper, we briefly touched motif discovery and protein classification tasks as use cases of peptide pair encoding representation. However, the application of this work is not limited to motif discovery or embedding training, and we expect this representation to be widely used in bioinformatics tasks as general purpose variable-length representation of protein sequences.

## Acknowledgements

Fruitful discussions with Hengameh Shams, Iddo Friedberg, and Ardavan Saeedi are gratefully acknowledged.

## Additional Information

### Author contributions

E.A. and M.R.K.M. conceived the project. E.A. implemented the method and wrote the main manuscript text. E.A, A.C.M, and M.R.K.M. contributed to the evaluations. All authors reviewed the manuscript.

### Competing interests

The authors declare no competing interests.

